# Neural network extrapolation to distant regions of the protein fitness landscape

**DOI:** 10.1101/2023.11.08.566287

**Authors:** Sarah A Fahlberg, Chase R Freschlin, Pete Heinzelman, Philip A Romero

**Affiliations:** Department of Biochemistry, University of Wisconsin–Madison, Madison, WI, USA; Department of Chemical & Biological Engineering, University of Wisconsin–Madison, Madison, WI, USA

## Abstract

Machine learning (ML) has transformed protein engineering by constructing models of the underlying sequence-function landscape to accelerate the discovery of new biomolecules. ML-guided protein design requires models, trained on local sequence-function information, to accurately predict distant fitness peaks. In this work, we evaluate neural networks’ capacity to extrapolate beyond their training data. We perform model-guided design using a panel of neural network architectures trained on protein G (GB1)-Immunoglobulin G (IgG) binding data and experimentally test thousands of GB1 designs to systematically evaluate the models’ extrapolation. We find each model architecture infers markedly different landscapes from the same data, which give rise to unique design preferences. We find simpler models excel in local extrapolation to design high fitness proteins, while more sophisticated convolutional models can venture deep into sequence space to design proteins that fold but are no longer functional. Our findings highlight how each architecture’s inductive biases prime them to learn different aspects of the protein fitness landscape.

## Introduction

Protein engineering can be envisioned as a search over the sequence-function landscape to discover new proteins with useful properties^1^. Machine learning (ML) accelerates the protein engineering process by integrating experimental data into predictive models to direct the landscape search^2^. These models improve the quality of protein variants tested and reduce the total number of experiments needed to engineer a protein^3^. ML-assisted protein engineering approaches have expedited engineering of diverse proteins such as viral capsids, improved fluorescent proteins, and highly efficient biocatalysts^4–7^.

Broad classes of machine learning and deep learning techniques approach protein design from different perspectives^8–11^. We focus on supervised learning, which learns from experimental sequence-function examples to predict new sequence configurations with some intended biochemical/biophysical properties. There are many different classes of models that each make different assumptions about the underlying landscape, influencing the way sequence-function relationships are learned^2^. For example, linear models assume additive contributions from individual mutations and are unable to capture epistatic effects. More sophisticated convolutional neural networks use convolving kernels to extract meaningful patterns from the input data that capture long range interactions and complex, non-linear functions^12^. Many studies have assessed the predictive performance of ML models on existing protein sequence-function datasets^11–16^, but there is little work that rigorously benchmarks performance in real-world protein design scenarios with experimental validation^5,7,17^. ML-guided protein design is inherently an extrapolation task that requires making predictions far beyond the training data, and evaluating models in this task is challenging due to the massive number of sequence configurations that must be searched and tested^18^.

In this paper, we evaluate different neural network architectures’ ability to extrapolate beyond training data for protein design. We develop a general protein design framework that uses an ML model to guide an *in silico* search over the sequence-function landscape. We use this approach with different model architectures to design thousands of protein G (GB1) variants that sample a massive sequence space far outside the model’s training regime. The different models prioritize distinct regions of the landscape and display unique preferences for the sequence positions mutated and types of amino acid substitutions. We then experimentally test the designs using a high-throughput yeast display assay that evaluates variant foldability and IgG binding. We find all models show the ability to extrapolate to 2.5-5x more mutations than the training data, but the design performance decreases sharply with further extrapolation. The simple fully connected neural networks showed the best performance for designing variants with improved binding relative to wildtype GB1. Intriguingly, we also found that the parameter-sharing convolutional models could design folded, but non-functional, proteins with sequence identity as low as 10% from wildtype, suggesting these models are capturing more fundamental biophysical properties related to protein folding. Our high-throughput screen identified multiple designs with improved binding relative to wildtype GB1 and previously designed variants. Our work provides a rigorous assessment of model architectures’ capacities for protein design and will support ongoing advances in ML-driven protein engineering.

### Extrapolating learned protein fitness landscapes

Protein sequence space is nearly infinite and experimental methods can only characterize a very localized and minuscule fraction of this space^1^. Machine learning (ML) models trained on sparse sequence-function data infer the full fitness landscape and can make predictions for previously unobserved protein sequences. These models can guide a search through sequence space to discover protein designs with high predicted fitness **(Fig. 1a)**. Many of these predictions extend far beyond the training data and thus are extrapolations on the fitness landscape. Although model performance is known to degrade as predictions are made further from the training regime, it remains unclear how far these models can extrapolate to design high-fitness sequences^5,7^.

**Figure 1:**
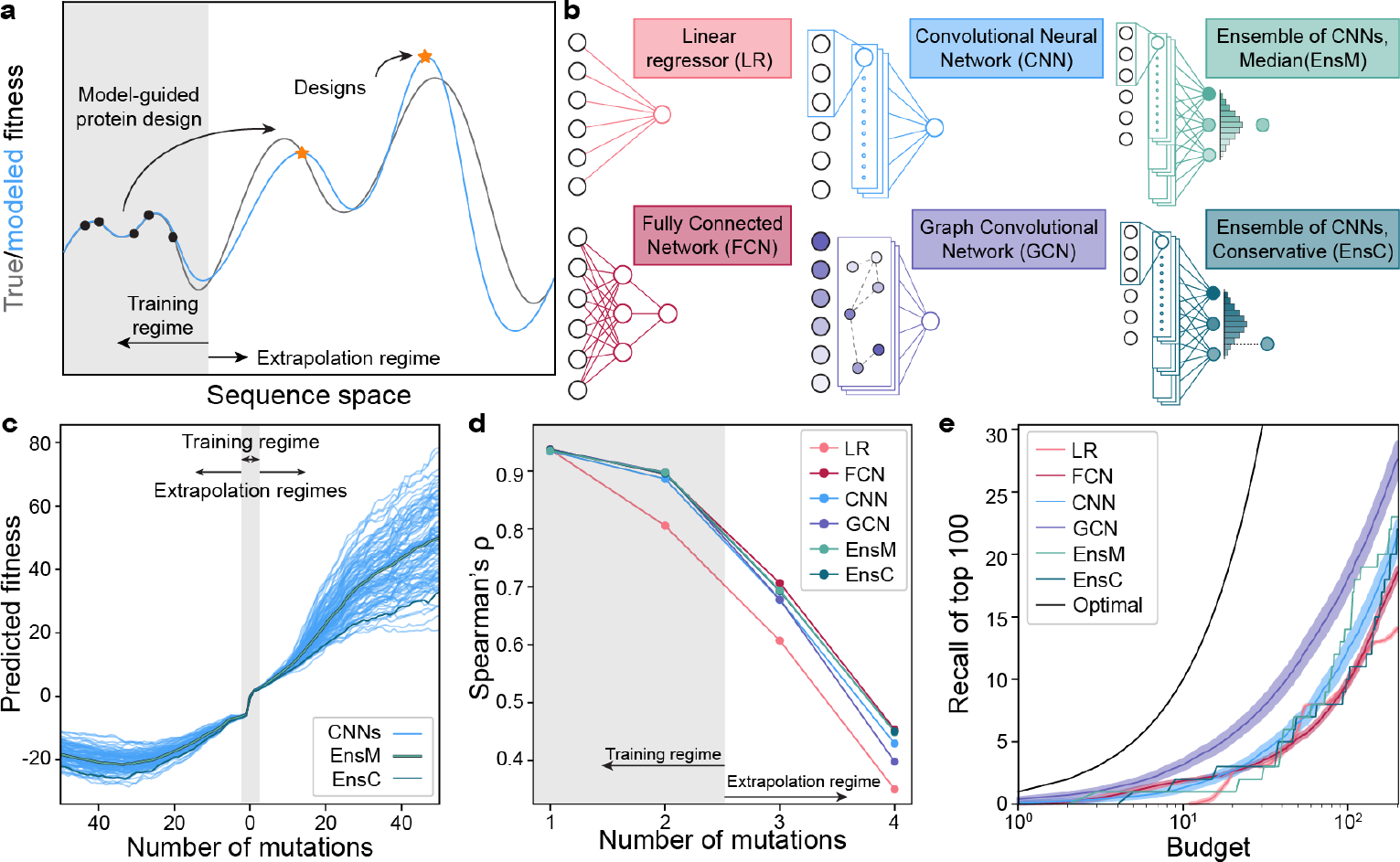
Extrapolation of sequence-function models. **(a)** Supervised sequence-function models are trained on experimental data and can make predictions across the fitness landscape. ML-guided protein design seeks to identify high fitness sequences and often involves model extrapolation beyond the training regime. **(b)** We tested five model architectures that capture distinct aspects of the underlying sequence-function landscape. **(c)** A collection of 100 CNN models and their divergence when predicting deep into sequence space along a mutational trajectory. The ensemble predictor EnsM is represents the median of the 100 CNNs, while EnsC is the 5^th^ percentile. **(d)** We trained models on GB1 single and double mutants and predicted the fitness of 1-, 2-, 3-, 4-mutants. The Spearman’s rank correlation was determined between the model’s predicted fitness and experimental fitness. **(e)** Model recall of the top 100 protein variants within a design budget. Recall represents the number of the true top 100 4-mutants that are present in a model’s top *N* predictions, where *N* is the design budget. Optimal represents a theoretical model that always predicts the top *N* proteins. Shading represents 95% confidence intervals across 100 individually trained models, excluding EnsM and EnsC.

The B1 binding domain of streptococcal protein G (GB1) is a small 8-kDa domain that binds to the mammalian Immunoglobin G (IgG) fragment crystallizable (Fc) region with nM affinity and to the fragment antigen-binding (Fab) region with *μ*M affinity^19,20^. We use GB1 as a model protein due to its extensively characterized IgG Fc-binding fitness landscape containing nearly all single and double mutations^20^. In previous work, we trained four different classes of supervised neural networks on GB1 double mutation data^12^. The networks included linear models that consider each residue position independently, fully connected networks (FCN) that can capture nonlinear behavior and epistasis, sequence-based convolutional neural networks (CNN) that share parameters across the protein sequence, and structure-based graph convolutional neural networks (GCN) that consider residues’ context within the protein structure (**Fig. 1b**). We previously evaluated these models’ predictive performance on held out double mutant data and demonstrated the feasibility of ML-guided protein design. However, it remains unclear how these models perform when extrapolating deeper into sequence space.

Neural networks contain hundreds of thousands to millions of parameters, many of which are not constrained by the training data^21^. The values of these unconstrained parameters are greatly influenced by the random initialization during training, and we hypothesized these parameters may lead to model divergence when predicting away from the training data regime. We trained 100 CNNs with the same model architecture, hyperparameters, and training data but with different random initializations of model parameters. We then evaluated each model’s predictions along a mutational pathway across the fitness landscape (**Fig. 1c**). All models show close agreement in the training data regime within two mutations of wildtype GB1, but their predictions increasingly diverged the further they moved from the training regime. It is also notable that the fitness predictions in the extrapolated regime have values so extreme they are unlikely to be valid. To overcome model variation arising from random parameter initialization, we implemented neural network ensemble predictors EnsM and EnsC that input a sequence into all 100 CNNs and return the median or lower 5^th^ percentile predictions for that sequence, respectively. We view EnsM as an “average” predictor, while EnsC is a conservative predictor because 95% of the models predict higher fitness values.

We evaluated the neural networks trained on single and double mutants on a separate GB1 data set containing nearly all combinations of mutations at four positions known to have high epistatic interactions^22^. In this dataset, single and double mutants are within the training regime, while prediction on the 3- and 4-mutants require model extrapolation. We found all models displayed significantly decreased predictive performance when extrapolating away from the training data (**Fig. 1d, Supplementary Fig. 1**). Although model accuracy drops dramatically in the extrapolated regime, the Spearman’s correlation remains significantly above 0, suggesting potential for model-guided design at or beyond 4 mutations. All the nonlinear neural network models performed similarly, while the linear model displayed notably lower performance, presumably due to its inability to model epistatic interactions between mutations.

We additionally tasked the models with identifying the most fit variants from the set of 121,174 possible 4-mutants, a task similar to machine-learning guided design. For a given testing budget *N*, each model ranks all 4-mutants and selects the top *N*. Recall is calculated as the proportion of the true top 100 that is represented in the model’s predicted top *N* sequences. For every budget tested, the GCN has the highest recall, indicating better extrapolation to identify high fitness variants (**Fig. 1e**). The FCN also has high recall with small design budgets but is quickly surpassed by the CNN.

### ML-guided protein design for deep exploration of the fitness landscape

Even a small protein like GB1 has over 10^70^ possible sequence configurations, and we must search deep into sequence space to fully evaluate ML models’ performance for protein design. We developed a large-scale protein design pipeline that uses simulated annealing (SA) to optimize a model over sequence space to identify high fitness peaks. The approach executes hundreds of independent design runs to broadly search the landscape, clusters the final designs to remove redundant or similar solutions, and then selects the most fit sequence from each cluster to provide a diverse sampling of sequences predicted to have high fitness (**Fig. 2a**). The number of clusters can be adjusted to match any downstream gene synthesis or experimental budgets.

**Figure 2.**
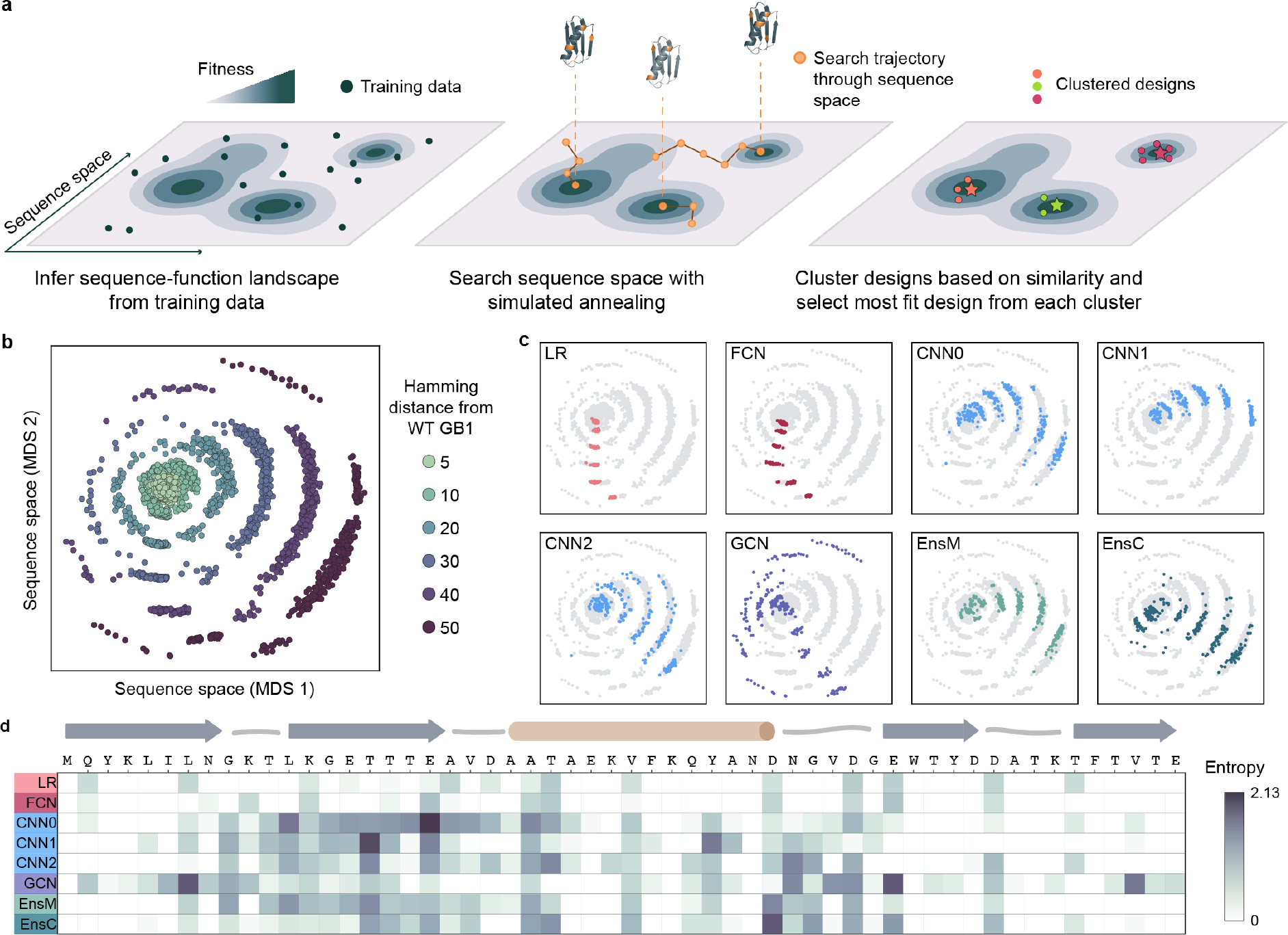
ML-guided fitness landscape exploration. **(a)** Supervised models infer the fitness landscape from sequence-function examples. We use simulated annealing (SA) to search through sequence space for designs with high predicted fitness. We perform hundreds of independent SA runs to broadly search sequence space, cluster designs to map distinct fitness peaks, and select the most fit sequence from each cluster. **(b)** We visualized all designs using multidimensional scaling (MDS) and found the designs occupy concentric rings emanating from wild-type GB1 with increasing number of mutations. **(c)** We colored the MDS visualization by model architecture and found individual models design sequences that occupy distinct regions of sequence space. **(d)** We calculated the sequence diversity across GB1 positions for the 10-mutant designs. We used Shannon entropy to quantify amino acid diversity at each position; low entropy indicates few amino acid options at a given site while high entropy indicates many amino acid options. The low entropy for the LR and FCN indicate that each model repeatedly proposes the same mutations at the same positions. The convolutional models propose sequences with more diversity spread across more positions, especially in regions of positive epistasis.

We applied our pipeline to design a diverse panel of GB1 variants testing different model architectures and spanning a range of extrapolation distances. We included eight models: LR, FCN, three CNNs with different initializations (CNN0, CNN1, CNN2), GCN, and the ensembles EnsM and EnsC. For each model, we designed variants at six extrapolation distances: 5, 10, 20, 30, 40, and 50 mutations from wild-type GB1. For each model-distance combination, we ran at least 500 design runs and clustered the designs into 41 clusters, to obtain 41 diverse sequences for each criterion. We visualized the design space using multi-dimensional scaling, which organizes sequences in a 2D space that attempts to preserve sequence interrelationships. We observe that sequences occupy concentric rings expanding outward from wildtype GB1, with each successive ring representing an increased mutation distance and the outermost ring corresponding to the 50 mutants (**Fig. 2b**). We observed notable differences in the sequences designed by each model, suggesting each architecture prioritizes distinct regions of the landscape (**Fig. 2c, Supplementary Fig. 2**). The LR and FCN designs occupy similar regions of sequence space and tended to display less variation within the 41 designs, suggesting a smooth inferred landscape structure with a major prominent peak. The designs from the three CNNs were distinctly different than the LR and FCN designs and displayed a high degree of variation across each model, highlighting how random parameter initialization can greatly influence model extrapolation. The GCN designs were by far the most diverse, occupying all directions in sequence space and having the highest average Hamming distance, reminiscent of a landscape with many distinct fitness peaks. At lower mutational distances, EnsM and EnsC produced designs that were more similar to that of LR and FCN, but in higher mutational regimes, their designs were more similar to designs produced by individual CNNs (**Supplementary Fig. 2**).

We further explored the unique mutational variation of each architecture’s designs by analyzing the amino acid diversity at each site (**Fig. 2d**). Generally, the LR and FCN target only a few positions in the protein sequence and tend to propose the same mutations, as indicated by the low entropy value. Every mutation proposed by the LR and FCN are individually beneficial (**Supplementary Fig. 3**). In contrast, the CNNs and GCN target a much broader region of the GB1 sequence, including known sites of positive epistasis around residues 9-16 and 33-40^20^. This suggests the CNN and GCN models can exploit epistasis to design sequences composed of synergistic mutations.

**Figure 3.**
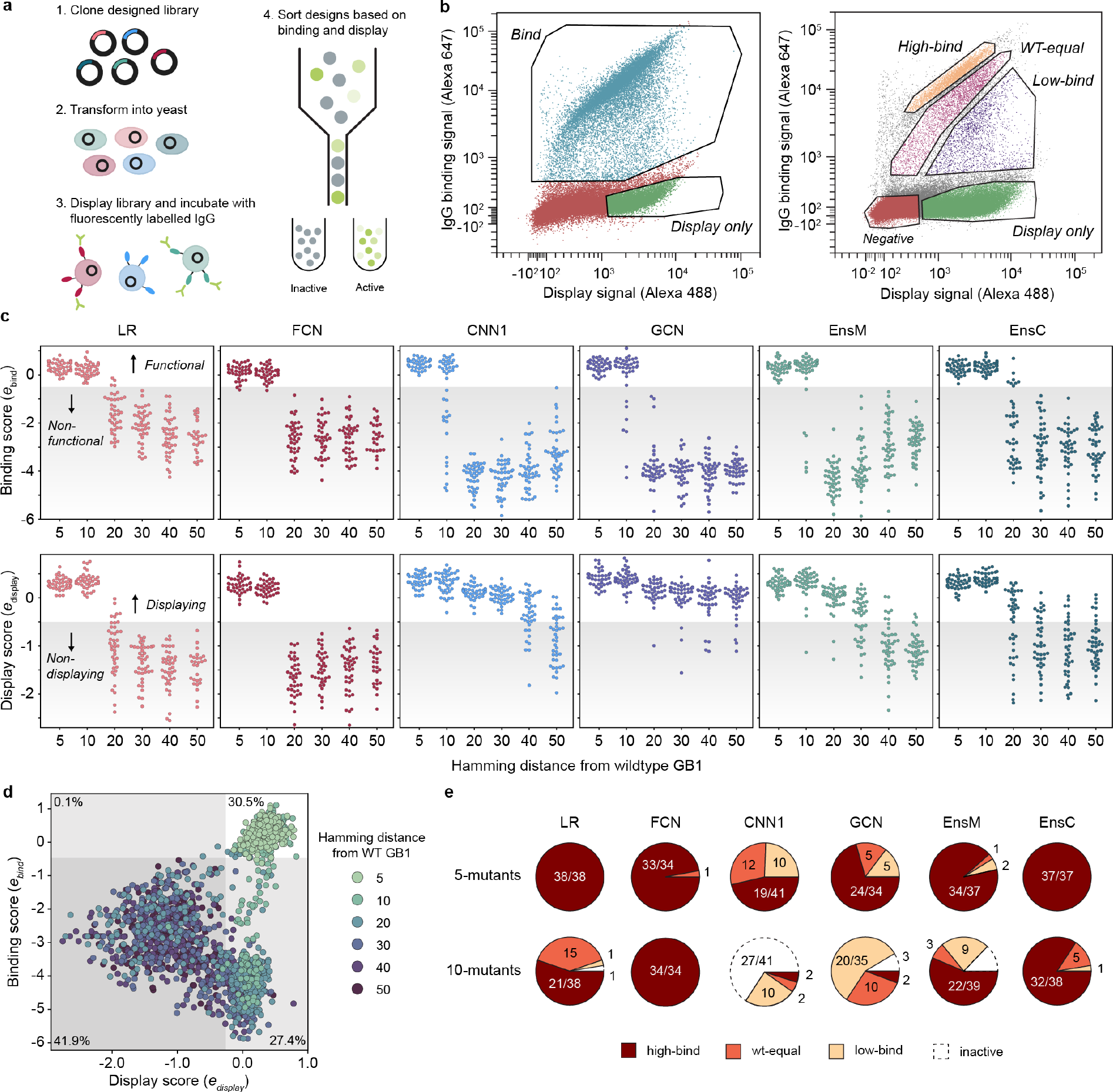
Experimental characterization of ML-designed GB1s. **(a)** An overview of our yeast surface display method to measure IgG-binding of GB1 designs. **(b)** FACS scatter plots and sorting gates for the library sorting experiments. In our first “qualitative” experiments, we sorted variants into “bind” and “display” populations. In our second “quantitative” experiment, the library was sorted into “display”, “low-bind”, “wt-equal” binding, or “high-bind” categories. **(c)** The binding and display scores for each model as a function of the Hamming distance from wildtype GB1. In the plot, each point corresponds to a single design and the shaded region specifies the threshold between functional/nonfunctional and displaying/non-displaying. **(d)** A scatter plot of display and binding scores for each design. The gray shaded regions specify the thresholds between functional/nonfunctional and displaying/non-displaying. The percentage of designs falling within each quadrant is specified in each quadrant’s corner. **(e)** The distribution of 5-mutant and 10-mutant designs categorized as high-bind, wt-equal, low-bind, or inactive from the quantitative experiment. Most designs beyond 10 mutations were inactive.

### Large-scale experimental characterization of ML designed GB1 variants

We used yeast surface display to characterize the expression and IgG binding of the designed GB1 variants. We sorted variants into “displaying” and “IgG binding” populations using fluorescence-activated cell sorting (FACS) (**Fig. 3a, b**) and sequenced these sorted populations to determine which variants fell into each bin. We used enrichment to devise *display* and *IgG binding* enrichment scores, *e*_*display*_ and *e*_*bind*_ respectively, for each variant. The display and binding enrichments show good reproducibly between experimental replicates and internal standards with varied nucleotide sequences but identical amino acid sequences, and the IgG binding enrichment correlates with the fitness value from the original deep mutational scanning study (**Supplementary Fig. 4**). The display score captures a combination of protein expression, trafficking, folding, and stability, while the IgG binding score is more directly related to IgG binding affinity. Additionally, display is a prerequisite for binding because the GB1 protein must reach the cell surface to interact with IgG.

**Figure 4:**
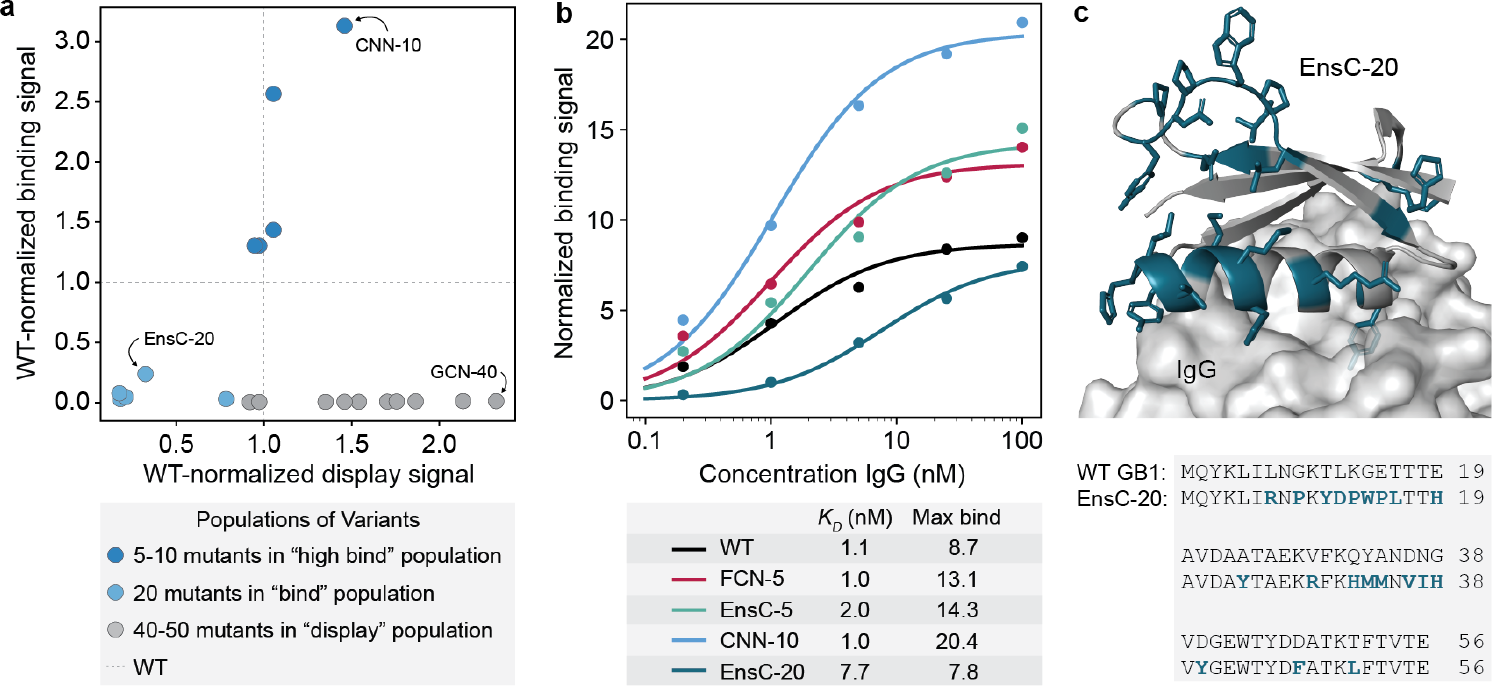
Validation of high-throughput yeast-display screen. **(a)** Clonal yeast display assay to verify designs’ display and IgG binding properties. The display and binding signals were normalized to wildtype GB1. **(b)** IgG binding curves for wildtype GB1 and designs FCN-5 (a 5 mutant designed by FCN), EnsC-5 (a 5 mutant designed by EnsC), CNN-10 (a 10 mutant designed by a CNN) and EnsC-20. We analyzed cells using flow cytometry and the normalized binding signal is the ratio of IgG binding to display level. We estimated the *K*_*D*_ and max binding signal parameters by fitting the data to the Hill equation. **(c)** AlphaFold2 predicted structure of the binding 20-mutant variant EnsC-20. The mutated residues are shown as sticks and highlighted in teal. IgG was included from the GB1 crystal structure (PDB: 1FCC).

We found all models were successful at designing functional GB1 variants that displayed and bound IgG (**Fig. 3c**). The design success rate was high at five and ten mutations and significantly decreased with further extrapolation from the training data regime. The simple LR and FCN models outperformed the more sophisticated CNN and GCN models for designing functional sequences that bind to IgG. Interestingly, the ensembles, which were composed of CNNs, showed design performance comparable to the LR/FCN models. This suggests the CNN random initialization process may result in models with subpar performance, but this effect can be marginalized by averaging over many models. CNN ensembles combine the strong generalization performance of parameter sharing models^12^ with the extrapolative design ability of simpler LR/FCN models.

The model design results for the display score were strikingly different than for IgG binding. While the LR and FCN design success decreased sharply with increasing extrapolation, the CNN and especially the GCN designed sequences that displayed out to 50 mutations. The picture becomes more complete when we jointly visualize the binding and display scores over extrapolation distances (**Fig. 3d**). The sequences can be broadly categorized into the four quadrants binds/doesn’t bind and displays/doesn’t display. The non-binding LR and FCN designs tended to not display, while the non-binding CNN and GCN designs displayed, suggesting two distinct modes of functional inactivation (**Supplementary Fig. 5 and Supplementary Fig. 6**).

We can contextualize these results if we consider that stable, folded proteins are more likely to display on the yeast surface, while unstable proteins are prone to proteolytic degradation^23^. In this case, the LR/FCN designs are inactive because they are unfolded, while the CNN/GCN designs are folded but may have a defective IgG-binding interface. The model architectures and their intrinsic inductive biases may explain the differences in each model’s design behavior. The LR/FCN models put an emphasis on specific sites and how those sites contribute to function. In contrast the CNN/GCN models have parameter sharing architectures that learn general patterns across the protein sequence. The site-focused designs from the LR/FCN models will maintain an intact IgG-binding interface, at the potential cost of losing protein stability. While the CNN/GCN models learn more general rules related to protein folding, but in the process, put less emphasis on the IgG binding interface.

We observed that some of the yeast display results may be explained by the presence of KEX2 protease sites in the designed proteins. KEX2 is located in the yeast Golgi and cleaves secreted proteins with exposed Lys/Arg-Arg sites. Unfolded proteins with KEX2 sites will be cleaved and not display, while folded proteins with KEX2 sites will typically display^24^. We found designs with 20 or more mutations are less likely to display if they have a KEX2 site (**Supplementary Fig. 7**). We also found the LR and FCN models had a higher likelihood of designing sequences with KEX2 sites (**Supplementary Fig. 8**).

We performed an additional yeast display experiment to resolve more quantitative differences in IgG binding by sorting variants into four bins corresponding to greater than WT binding (high-bind), WT-like binding (WT-equal), lower than WT binding (low-bind), and no IgG binding (display only) (**Fig. 3b**). Our results reveal that all model architectures succeed in designing functional GB1 variants with five and ten mutations (**Fig. 3e**). When considering the individual models, the LR and FCN significantly out-perform the CNNs and GCN with nearly 100% of designs falling in the high-bind category for both five and ten mutations. While the GCN and CNNs design some functional ten-mutants, few designs are high-binding. In contrast to the individual CNNs, the ensembles are as successful as the LR and FCN in designing high-bind GB1 variants, suggesting that ensembles of models confer some benefit when extrapolating beyond the training regime.

### ML designed GB1s show improved display and IgG binding

We used the high-throughput screening data to identify 20 interesting designs for more detailed functional and structural analysis **(Supplementary Table 2)**. We chose ten designs with 40-50 mutations that displayed but did not show IgG binding and we chose another ten designs with 5-20 mutations that showed IgG binding. We quantitatively evaluated display and IgG binding for each of these 20 designs in a clonal yeast display assay **(Fig. 4a)**. 9/10 of the “display” designs displayed on the yeast surface equal to or greater than wildtype GB1, including a 40-mutant designed by the GCN that displayed over 2x higher than wildtype. This variant, called GCN-40, shares less than 30 % sequence identity with wildtype GB1 and is predicted to have some similar structural elements, but with a completely new helical bundle fold **(Supplementary Fig. 9)**.

5/5 of the “high binding” designs with 5-10 mutations showed IgG binding greater than wildtype; however, the 5 “binding” designs with 20 mutations showed diminished binding. We performed IgG titrations to assess the binding curves for several designs **(Fig. 4b)**. While the designs do not have significantly different *K*_*D*_s from wildtype (**Fig. 5c, d**), the maximum binding signals are significantly larger for the designs, suggesting our designs have decreased *k*_*off*_ rates in our yeast display assay. We hypothesize the initial GB1 training data mRNA display experiments^20^ may have captured *k*_*off*_ effects rather than equilibrium *K*_*D*_ measurements and thus resulted in trained models that design GB1 variants with improved binding kinetics but not necessarily thermodynamics. The most distant functional design was a 20-mutant designed by EnsC, termed EnsC-20, that showed significant IgG binding, although much weaker than wildtype. Many of the mutations in EnsC-20 are at the IgG interface, while others, including three mutations to proline, were on the distal side of the protein, likely inducing structural changes that could affect binding.

## Discussion

Protein engineering has broad applications across biocatalysts, biomanufacturing, agriculture, and human health, but engineering proteins is slow and requires expensive rounds of experimental characterization to search the fitness landscape. Machine learning accelerates the protein engineering process by inferring the underlying landscape structure and reducing the experimental burden required to explore protein sequence space. In this work, we assessed different supervised machine learning architectures’ ability to extrapolate beyond the training data for protein design. Adapting previously developed models trained on a large GB1 sequence-function dataset, we performed an ML-guided search to design sequences predicted to have high fitness. We found each model had markedly different perceptions of the underlying landscape that gave rise to unique preferences for the designed sequences. All models showed strong potential to extrapolate to 2.5-5x more mutations than represented in the training data, but the simpler fully connected architecture performed the best for designing functional and highly fit proteins.

Our work rigorously evaluates a common protein engineering setting wherein a supervised sequence-function model is trained on a local fitness landscape and tasked with extrapolating to new regions of sequence space^5,6,11,24,25^. These models become increasingly inaccurate further from the training data due to the challenge of generalizing local sequence rules to the global landscape, which is exacerbated by factors like landscape ruggedness, data sparsity, and magnitude of epistasis. Models that effectively capture epistasis and that can learn generalizable biophysical concepts will be more capable of extrapolating on the fitness landscape. Assessing a model’s extrapolation potential requires more than simply holding out some subset of variants from a localized and fixed data set. Indeed, our analysis found CNN/GCN models that perform well in predicting 4-mutants (**Fig. 1**) tend to underperform when extrapolating to 5, 10, and 20 mutations (**Fig. 3**). These findings highlight differences between local and global landscape structures and the need to design and experimentally test sequences to evaluate model performance.

A given model’s extrapolation ability will depend on the size and quality of the training data, the model’s inductive biases and its capacity to learn sequence-function relationships, and the ruggedness of the optimization surface, which can limit search methods’ ability to identify top solutions. Despite simulated annealing’s (SA) ability to bypass local optima, we observed notable differences in the spread of each model’s designs (**Fig. 2c**), suggesting varied degrees of ruggedness in the underlying optimization surface. The SA trajectories for the LR designs tended to converge toward the same sequences, as expected for a smooth landscape with a single global peak. In contrast, the GCN designs were broadly distributed across sequence space, indicating each SA trajectory had arrived at distinct local optima due to the difficulty in traversing a highly rugged optimization surface.

We found the LR and FCN models strongly outperformed the convolutional models in designing functional 5- and 10-mutants that bound IgG tighter than wildtype GB1. These models have a simple architecture where each position in the amino acid sequence contributes to function and highlights the relative simplicity of the local landscape for a given property. Additive mutational effects have been known and leveraged in protein engineering for decades^26,27^, and the LR and FCN’s inductive biases are set up to capture these simple relationships. FCNs have the additional advantage that they can learn epistatic relationships between residues, which resulted in increased performance relative to LR. The primary distinguishing feature of the convolutional models is their parameter sharing architecture that learns filters that are applied across the protein sequence/structure. Convolutional models have the capacity to learn more general relationships, but their inductive biases are not primed to capture the simpler additive and epistatic relationships that dominate local landscapes.

We interpret our results primarily through the lens of protein engineering where our goal is to design GB1 variants with improved binding to IgG. But we found the CNN, and especially the structure-based GCN, were highly effective at designing GB1s with up to 50 mutations that display on the yeast surface and presumably fold, but do not bind IgG. It’s worth noting that this phenomenon doesn’t necessarily indicate subpar model performance, and instead we postulate the parameter sharing architectures may have focused more on learning general rules of protein folding and, in the process, ignored IgG binding activity. While the original sequence-function training data was based on IgG binding, this experimental mapping is dominated by folding effects because a majority of a protein’s residues are involved with maintaining structural stability, while relatively few are directly involved with binding. In other words, a mutation can inactivate a protein due to disrupted folding or disrupted binding, but the disrupted folding mechanism is statistically much more likely. It’s well known that most mutations destabilize proteins^28^ and this observation was noted in the original GB1 deep mutational scanning manuscript^20^. The combination of data that’s dominated by folding effects and the convolutional models’ inductive bias to recognize patterns led to models that learned more general rules of protein folding.

We found ensembles of CNNs outperform individual CNNs in designing high fitness variants. Ensembles highlight model variation arising from random parameter initialization and can identify regions of sequence space that are poorly understood by the model. Being able to identify where a model is not confident is critical for robust design. Model confidence can be estimated by inputting a single sequence into an ensemble of models and evaluating the agreement in prediction across the models^29^. If the models tend to agree, we can be more confident in the prediction, while if the ensembles’ predictions are highly variable, then the random parameter initialization may be influencing the models’ prediction. Evaluating this consensus between models is important for neural network models that may have millions or more internal parameters that are not fully determined by the data. Other work has used model uncertainty to guide designs toward regions of sequence space that are generally more confident^21,30–32^. For our EnsC predictor, we do not restrict design space based on uncertainty, but rather use the ensemble to make a conservative fitness prediction where 95% of the models agree the fitness is above a certain value. Designing sequences that maximize EnsC result in designs that most models agree have high fitness. We found the EnsC design strategy was the best overall with similar performance to LR and FCN for designing highly fit 5- and 10-mutants and some ability to design functional sequences with 20 mutations.

Protein function is multifaceted; a single output property such as binding may be influenced by mutations that affect structure, the binding interface, pH sensitivity, or protein expression. Similar to the adage ‘you get what you screen for,’ in ML-guided protein engineering, you get what you train on. It is difficult to know a priori the mechanisms though which individual mutations affect fitness. We found different model architectures are more primed to learn distinct underlying mechanisms. This naturally leads to the idea that these differing architectures could be used together to design functional proteins more robustly. A collection of model architectures, each with their own inductive biases, could be used together in an ensemble, similar to EnsC. Ensembles of different model classes have been successfully used for protein and metabolic engineering^7,33^. Another design strategy could aim to jointly optimize multiple models simultaneously using multi-objective criteria^34^.

Deep learning and artificial intelligence are revolutionizing the fields of protein science and engineering. These tools can process, learn from, and make sense of large quantities of data to decode the complex inner workings of proteins with a scale and resolution beyond human comprehension. Continued advances will help realize the potential of protein design to address society’s most pressing problems and future global challenges.

## Methods

### Neural network model training

We used the Olson et al. GB1-IgG binding dataset^20^ and the Gelman et al. model architectures^12^ with the same parameter initializations, train-validation-test splits, and hyperparameters. We additionally trained 100 models of each architecture with different random initializations to assess the effects of parameter initialization and for ensemble learning. We constructed two CNN ensemble predictors: a median predictor (EnsM) that inputs a sequence into 100 CNN models and outputs the median fitness prediction, and a conservative predictor (EnsC) that inputs a sequence into 100 CNN models and outputs the 5^th^ percentile fitness prediction.

### Evaluation of model extrapolation on 3- and 4-mutant fitness landscapes

We evaluated the models’ ability to extrapolate to 3- and 4-mutants using the four-site combinatorial GB1 data set from Wu et al.^22^ This dataset consists of a complete 4-site combinatorial saturation mutagenesis GB1 library screened for binding to IgG. The models were trained on single and double mutant data from Olson et al.^20^ and were used to predict the fitness values of the 3- and 4-mutants. We also tested the ability of the models to identify the 4-mutants with the highest fitness values. The model recall was calculated by giving a model a design budget *n*, having the model rank all 121,174 characterized 4-mutants and select the top *n*, and then determining what percentage of the true top 100 variants were in this top *n* set. An optimal model would achieve 100% recall with a design budget *n* = 100.

### Model-guided protein design

We designed GB1 variants with eight different models at 5, 10, 20, 30, 40, and 50 mutations from wildtype GB1 (**Supplementary Table 1**). For each model-distance combination, we designed 41 diverse GB1 variants with high predicted fitness. For a given model-distance combination, we used simulated annealing (SA) to maximize the model’s predicted fitness constraining the number of mutations allowed in the designed protein. To ensure the number of mutations in a design were fixed, we exchanged random mutations. For example, when designing a 5-mutant, we might randomly choose two current mutations, mutate those back to the wildtype amino acids, and then choose two new random mutations to keep the total mutations fixed at five. At each step of simulated annealing, one or more mutations (sampled from a λ = 1 Poisson distribution) were randomly exchanged and predicted fitness of the given model was evaluated. Mutations that improved predicted fitness were automatically accepted while all other mutations were accepted with probability *e*^Δ*Fitness*/*T*^ where *T* is decreased along a logarithmic temperature gradient ranging from 10^3^ to 10^-5^ over 15,000 to 50,000 steps. We performed 500 independent simulated annealing runs for each model-distance combination, which typically resulted in more than 41 unique candidate sequences. For select model-distance combinations, 50,000 SA steps converged on fewer than 41 candidate sequences, in which case, we decreased the number of SA steps until more than 41 unique candidate sequences were designed. The number of SA steps for these specific categories were: (LR 5 mutants) and (FCN 5 mutants) = 10,000 steps and (LR-10 mutants), (FCN-10 mutants), (EnsM-5 mutants), and (EnsC-5 mutants) = 25,000 steps. All sequence designs were run in parallel using high-throughput computing^35^. To select a diverse and representative set of designs, we performed K-means clustering on the 500 independent designs using 41 clusters and selected the variant from each cluster with the highest predicted fitness for experimental characterization. This resulted in 41 unique GB1 designs for 8 models at 6 distances, for a total of 1968 designs.

### High-throughput characterization of GB1 designs via yeast surface display

We codon optimized our GB1 designs for expression in *S. Cerevisiae* using GenSmart Codon Optimization (GenScript) and excluded the BsaI, NheI, BamHI restriction enzyme sites. We also identified 25 sequences from the training data with a broad range of fitness values to correlate our fitness measurements with the original data from Olson et al.^20^. Each of these variants were designed with two different synonymous codon sequences to provide internal controls to ensure reproducibility of our fitness measurements. The designed genes and control sequences were synthesized as an oligonucleotide pool by Twist Biosciences with flanking sequences to allow PCR amplification and downstream cloning.

We amplified the oligonucleotide pool using either Phusion Hot Start Flex 2X Master Mix (New England Biolabs) or KAPA HiFi HotStart ReadyMix (Roche), cloned the gene library into the yeast surface display vector pCTCON2 (provided by Dane Wittrup, MIT)^36^, transformed into 10G supreme electrocompetent *E. coli* (Lucigen), and harvested the plasmid DNA using the Qiaprep Spin miniprep kit (Qiagen). We then transformed the GB1 library into yeast display *Saccharomyces cerevisiae* strain EBY100 made competent using the Zymo Research Frozen EZ Yeast Transformation II kit. We grew the transformed library in pH 4.5 Sabouraud Dextrose Casamino Acid media (SDCAA: Components per liter - 20 g dextrose, 6.7 grams yeast nitrogen base (VWR Scientific), 5 g Casamino Acids (VWR), 10.4 g sodium citrate, 7.4 g citric acid monohydrate) at 30°C and 250 rpm for two days. We plated an aliquot of the transformant pool on SD -Trp agar to quantify the number of library transformants.

We analyzed and sorted the GB1 library using florescence-activated cell sorting (FACS). We induced the library expression in SGCAA media overnight, harvested approximately 3e6 yeast cells by centrifugation, washed once in pH 7.4 Phosphate Buffered Saline (PBS) containing 0.2% (w/v) bovine serum albumin (BSA), and incubated overnight at 4 °C on a tube rotator at 18 rpm in 800 μL of PBS/0.2% BSA containing 15 nM human IgG1 (BioLegend) that had been conjugated with Alexa647 using NHS chemistry (Molecular Probes) and 3 μg/mL anti-*myc* IgY (Aves Labs) conjugated with Alexa488 using NHS chemistry. Following the overnight incubation, we washed the yeast in PBS/0.2% BSA and resuspended in ice cold PBS for FACS. We performed FACS using a FACS Aria II (Becton Dickinson) and analyzed yeast cells for display at 488 nm and IgG binding at 647 nm. Our “qualitative” experiments sorted cells into “display only” and “bind” populations, while our “quantitative” experiments sorted cells into “display only,” “low bind,” “WT-like bind,” and “high bind” populations.

We recovered the sorted yeast populations, as well as the initial unsorted library, and grew them in SDCAA media overnight. The following morning, we expanded the cultures into SDCAA media at an optical density of 0.1, grew them until reaching density of ∼1.0, harvested and centrifuged the cultures, and extracted the plasmids using the Zymo Research Yeast Plasmid Miniprep II kit. We transformed the extracted plasmid DNA into 10G supreme electrocompetent *E. coli*, cultured overnight in LB + carbenecillin media shaking at 250 rpm at 30°C, and harvested the plasmid DNA using the Qiaprep Spin miniprep kit (Qiagen). We cut out the GB1 gene insert using XhoI and PstI restriction enzymes, excised the corresponding band using agarose gel extraction, and purified using the Zymo Research gel extraction kit. The UW-Madison Biotechnology Center DNA sequencing core prepared a sequencing library using the NEBNext Ultra II kit and sequences the samples using an Illumina NovaSeq6000 with 2x150 bp reads.

### Illumina data processing and analysis

We aligned forward and reverse Illumina reads to wildtype GB1, using a predetermined offset, and merged the two reads by selecting the base with the higher quality score in overlapping regions. A design’s count was equal to the number of sequencing reads that exactly matched the designed nucleotide sequence and we filtered out any designs if they had fewer than 10 counts in the unsorted population.

The initial experiments consisted of two replicates each of unsorted (u), display only (d), and binding (b) populations. For each population, we divided by the total number of reads to obtain the proportion of each design. We refer to *<u>p</u>*_*<u>u,i</u>*_, *<u>p</u>*_*<u>d,i</u>*_, and *<u>p</u>*_*<u>b,i</u>*_ as the proportion of design *i* in the unsorted, display only, and binding populations, respectively. We calculated a binding score as the enrichment of a design in the binding population relative to the display only population, where *<u>p</u>*_*<u>b,wt</u>*_ and *<u>p</u>*<u>_*<u>d,wt</u>*_</u> are the proportion of wildtype GB1 in the binding and display only populations, respectively (**Equation 1**). We also calculated a display score as the enrichment of a design in the full displaying population (b+d) relative to the unsorted population (**Equation 2**). Here, the 0.6 and 0.4 correspond the relative proportion of the binding and display only populations from the FACS experiment. The numerator estimates the full displaying population from the binding and display only populations.

For the quantitative screening of designs, we obtained unsorted (u), display only (d), low binding (l), wildtype-like binding (w), and high binding (h) populations. For each population we calculated the enrichment relative to the unsorted population where *x* corresponds to any sorted population l, w, h (**Equation 3**). Each sequence was categorized into display only, low binding, wildtype-like binding, and high binding based on thresholds determined by calibration sequences with known fitness values (**Supplementary Fig. 10**). All data analysis can be found Jupyter notebooks in the supplementary material.

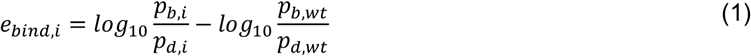

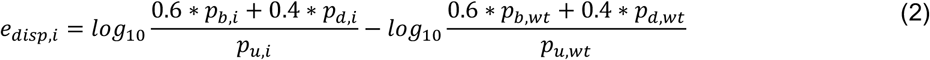

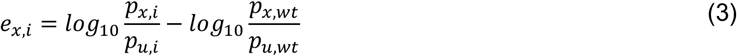

### Characterization of individual designs for display and binding

We identified twenty designs to test more thoroughly for binding and display in clonal yeast display assays (**Supplementary Table 2**). We chose five 5- and 10-mutants with high binding across all replicates, five 20-mutants with some binding activity across all replicates, and ten 40- and 50-mutants with high levels of display across all replicates. We resynthesized their DNA sequences as individual gene fragments (Twist Biosciences) and cloned them into the same yeast display vector used for library screening. We transformed the plasmid DNA into EBY100 made competent using the Zymo Research Frozen EZ Yeast Transformation II kit and grew the transformants on synthetic dropout (SD) -Trp (MP Biomedicals) agar plates for two days at 30°C. After two days, we picked individual colonies into 4 mL of pH 4.5 Sabouraud Dextrose Casamino Acid media (SDCAA: Components per liter -20 g dextrose, 6.7 grams yeast nitrogen base (VWR Scientific), 5 g Casamino Acids (VWR), 10.4 g sodium citrate, 7.4 g citric acid monohydrate) and grew them overnight at 30 °C and 250 rpm. We induced GB1 display in a 5 mL pH 7.4 SGCAA culture started at an optical density as measured at 600 nm of 0.5 and shaken overnight at 250 rpm and 20 °C.

For IgG binding titrations, we harvested approximately 2*10^5^ induced yeast cells for each titration data point, washed once in pH 7.4 Phosphate Buffered Saline (PBS) containing 0.2% (w/v) bovine serum albumin (BSA), and incubated for three hours at 4°C on a tube rotator at 18 rpm in between 100 μL and 1 mL of PBS/0.2% BSA containing various concentrations of Alexa647 human IgG1 and 3 μg/mL Alexa 488 anti-*myc* IgY. We varied the volumes of Alexa647 IgG-containing incubation solution to prevent ligand depletion from occurring at the lowest IgG concentrations. Following incubation, we washed the yeast once in PBS/0.2% BSA and resuspended in ice cold PBS for flow cytometric analysis. We analyzed the samples using a Fortessa analyzer (Becton Dickinson).

## Acknowledgements

We thank the University of Wisconsin Carbone Cancer Center Flow Cytometry Laboratory, supported by P30CA14520, 1S10OD018202-01, and 1S10OD018202-01, and the University of Wisconsin-Madison Next Generation Sequencing Core for assistance with experimental characterization of designs. This research was supported by the United States National Institutes of Health (R35GM119854, T32HG002760), the UW-Madison Biochemistry Undergraduate Summer Research Award, the UW-Madison Hilldale Undergraduate Research Award, and the UW-Madison Biochemistry Mary Shine Peterson Undergraduate Award.

