## Supplementary figures for "Neural network extrapolation to distant regions of the protein fitness landscape"

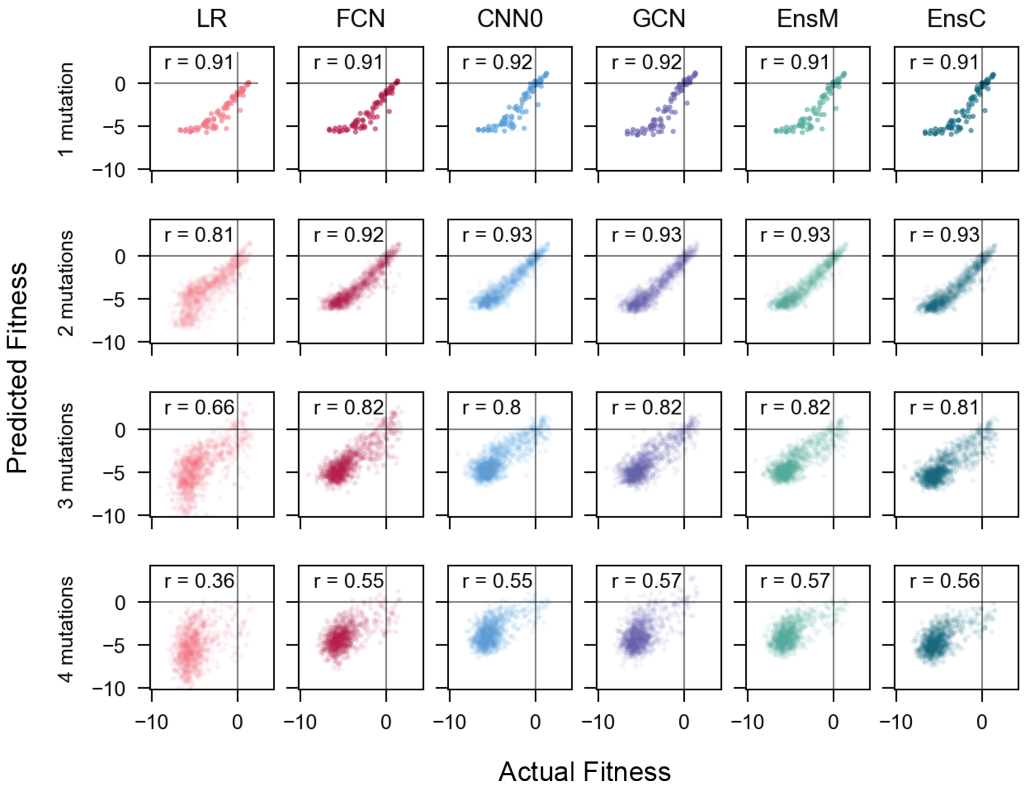


**Figure S1. Correlation between actual and predicted fitness of 1-4 mutant GB1s.** We trained models on single and double mutant GB1 fitness data from Olson et al.[1] and used each model to predict the fitness of all single, double, triple, and quadruple mutants from a 4-site GB1 combinatorial library from Wu et al.[2].

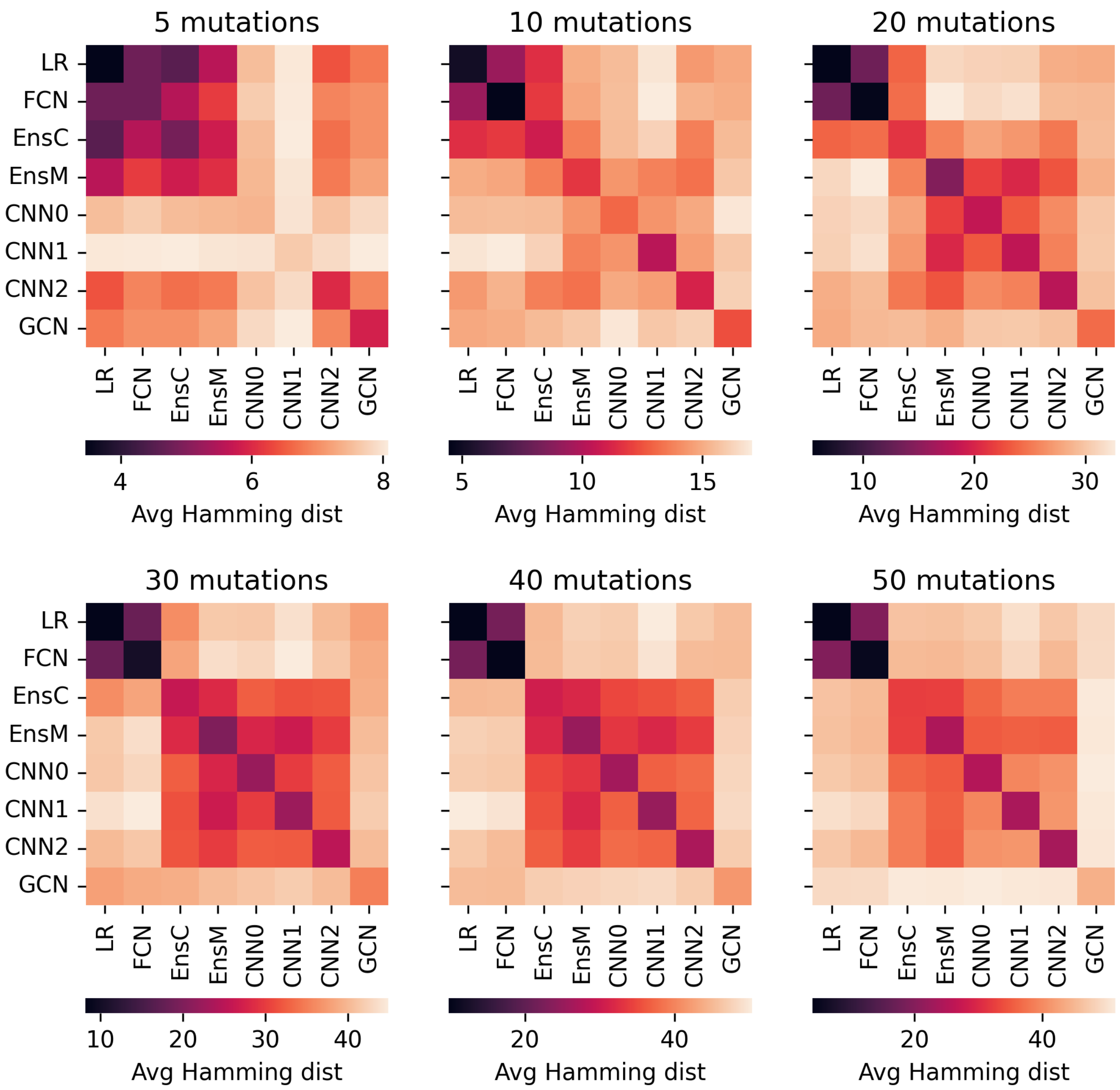


**Figure S2. Designs produced by each model show distinct similarity patterns across mutational distances.** We calculated the average pairwise Hamming distance between all designs from each pair of models and performed hierarchical clustering to group design strategies based on their similarity.

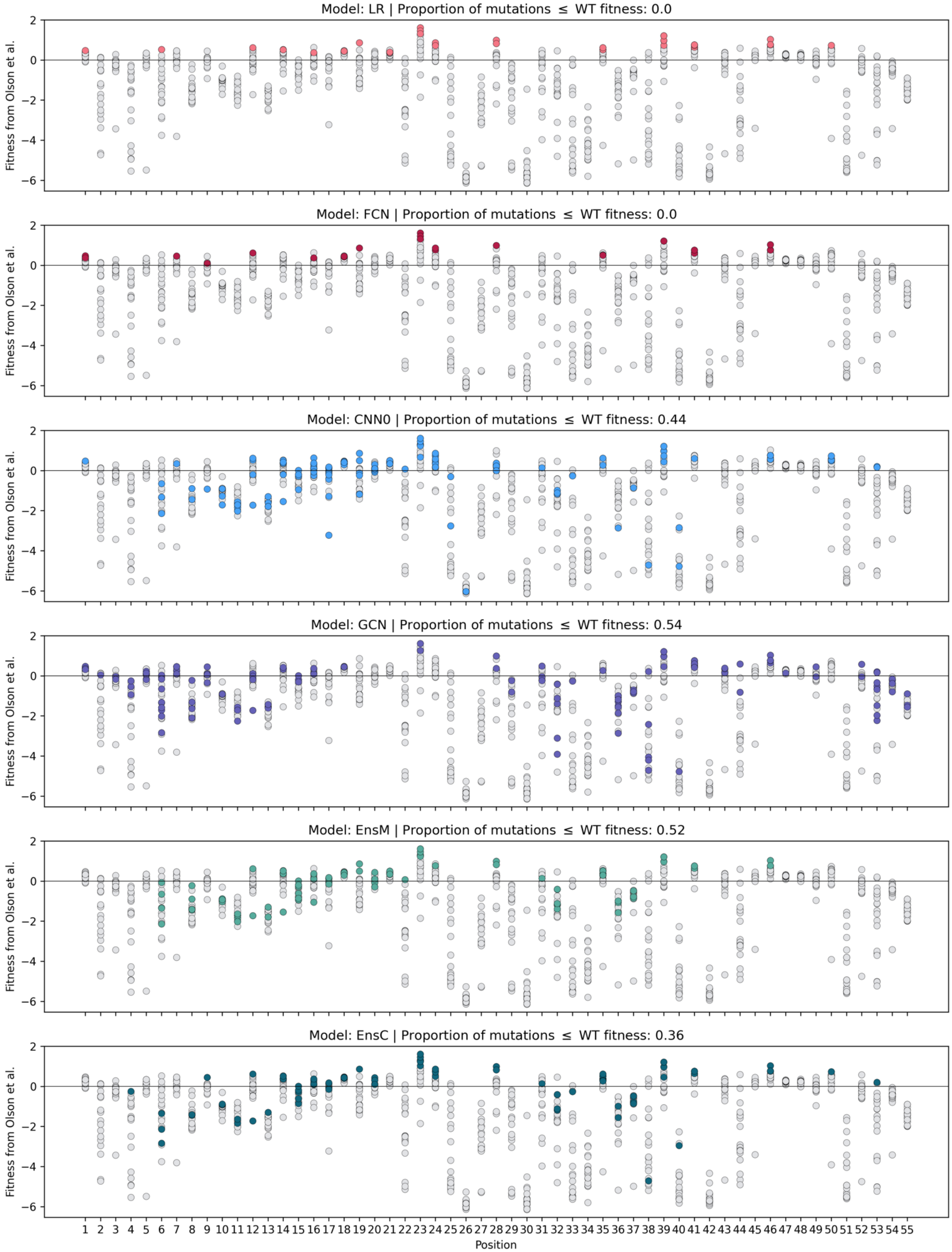


**Figure S3. Experimental values for all single mutations in designs with 10 mutations.** We show the original enrichment scores from Olson et al.[1] for all single mutants in grey. For each panel, we color the mutations found in each model's 10-mutation designs, highlighting the types of mutations chosen by each model.

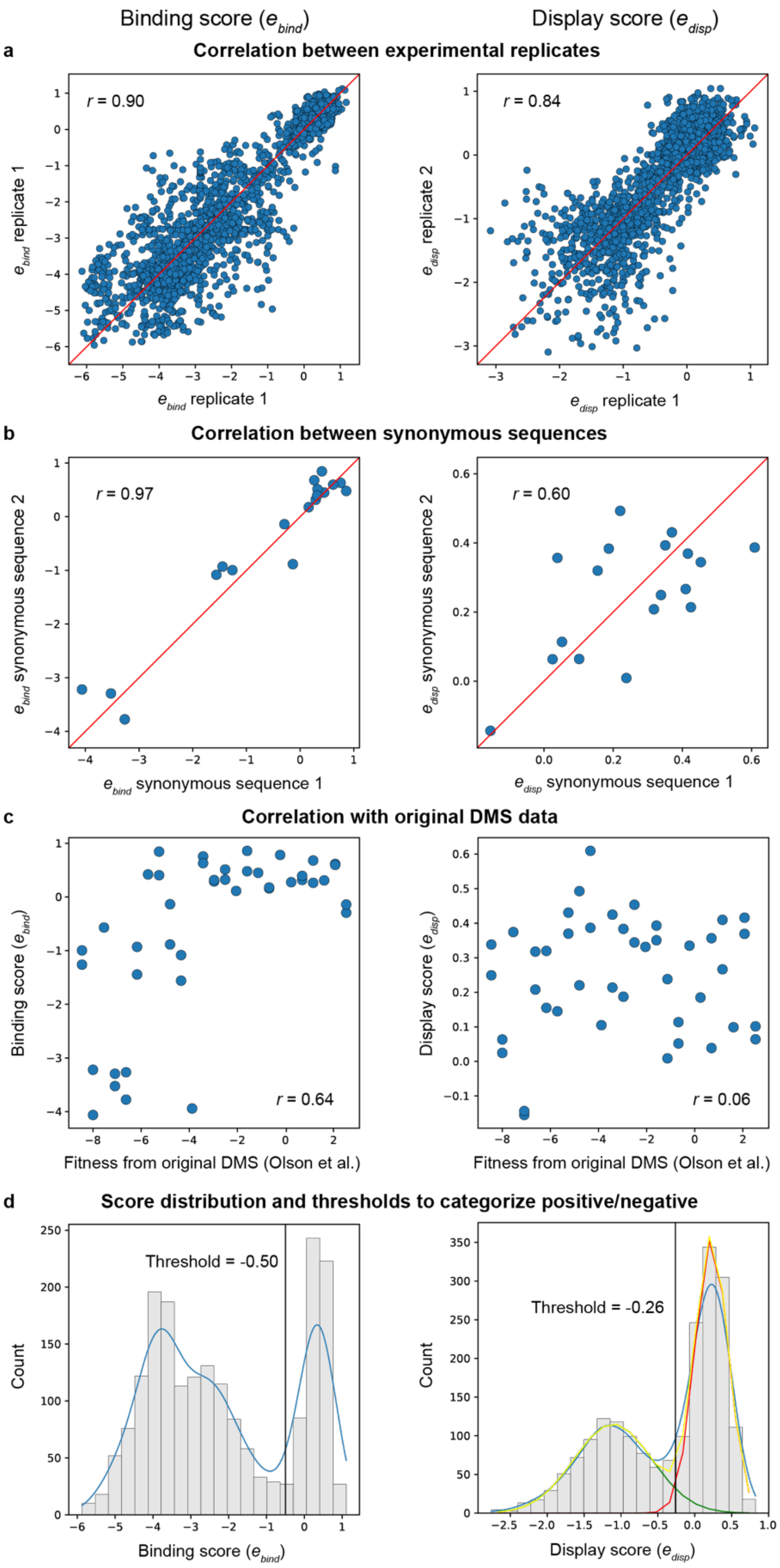


**Figure S4. Reproducibility of binding and display enrichment scores.** (a) The $e_{bind}$ and $e_{disp}$ scores show strong correlation between two independent experimental replicates. All subsequent analysis in the paper was performed on the average $e_{bind}$ and $e_{disp}$ between these two replicates. (b) In our experiments we included 19 synonymous sequence pairs with the same amino acid sequence but different nucleotide sequences that could be distinguished in the Illumina sequencing. The $e_{bind}$ and $e_{disp}$ scores for these synonymous sequence pairs show strong correlation indicating the assay is reliably measuring the protein's fitness. (c) In our experiment we also included 25 calibration sequences from the original deep mutational scanning (DMS) dataset to use as a reference. Our $e_{bind}$ score shows a moderate correlation with the DMS fitness, while $e_{disp}$ shows no correlation. (d) We manually categorized sequences as binding/not binding based on the distribution of $e_{bind}$ scores and manually setting a threshold to separate the two modes. We manually categorized sequences as displaying/not displaying based on the distribution of $e_{disp}$ scores, fitting a bi-Gaussian distribution, and identifying where the two Gaussians' densities cross.

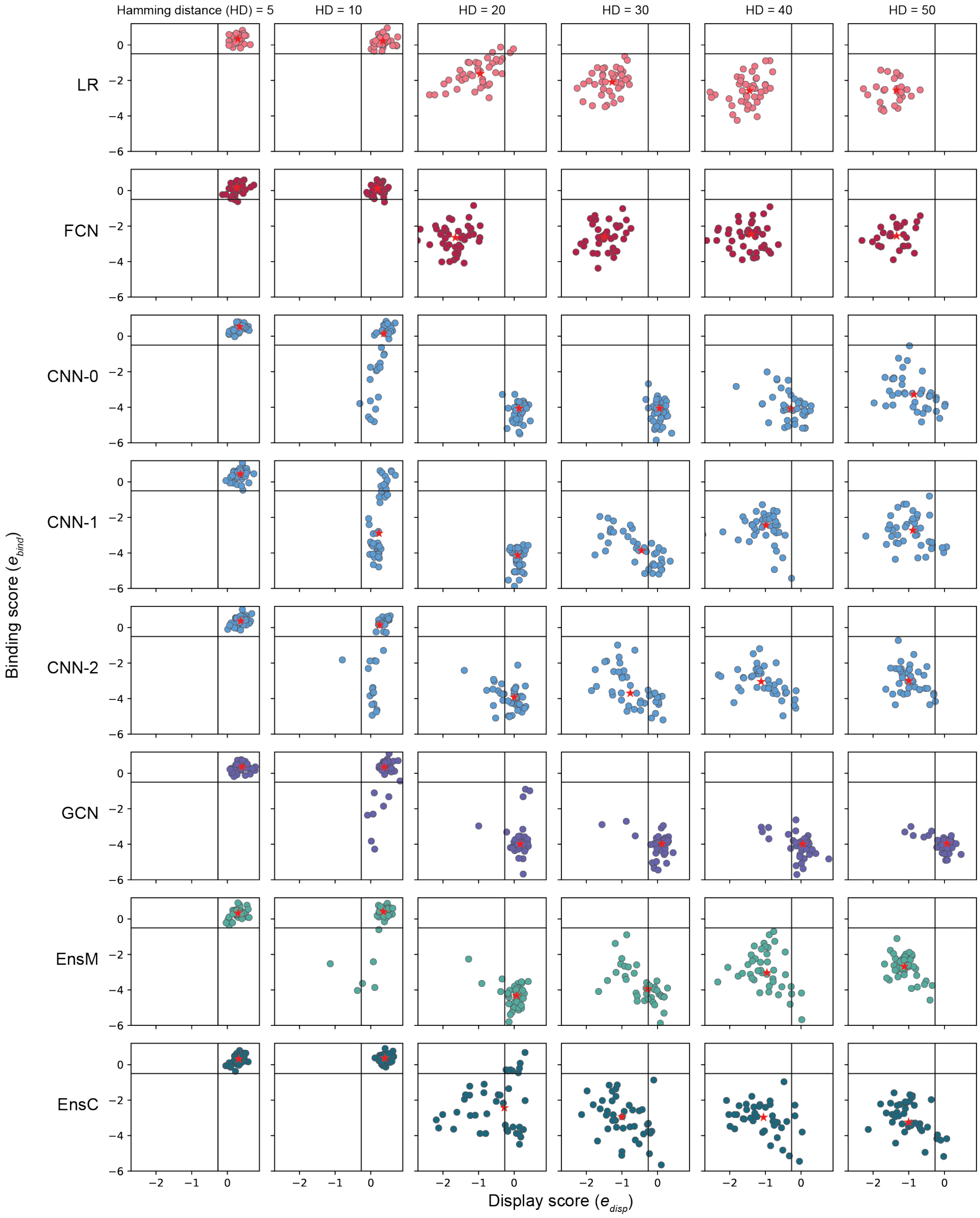


**Figure S5. Distribution of binding and display scores of designs for each model at each mutational distance.** Each design is shown as a colored point, plotted by its $e_{bind}$ and $e_{disp}$ scores. The median $e_{bind}$ and $e_{disp}$ score for each model at each mutational distance is shown as a red star.

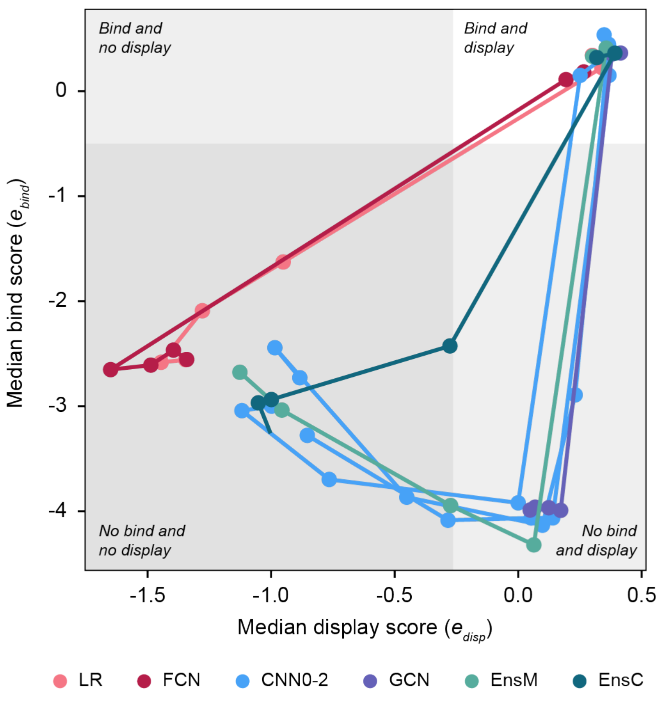


**Figure S6. Median $e_{bind}$ and $e_{disp}$ score trajectory for increasing mutational distance.** We calculated the median $e_{bind}$ and $e_{disp}$ scores for each model-mutational distance combination and plotted the trajectory with increasing mutational distance for each model. We overlay the $e_{bind}$ and $e_{disp}$ thresholds for bind/no bind and display/no display to separate the space into quadrants.

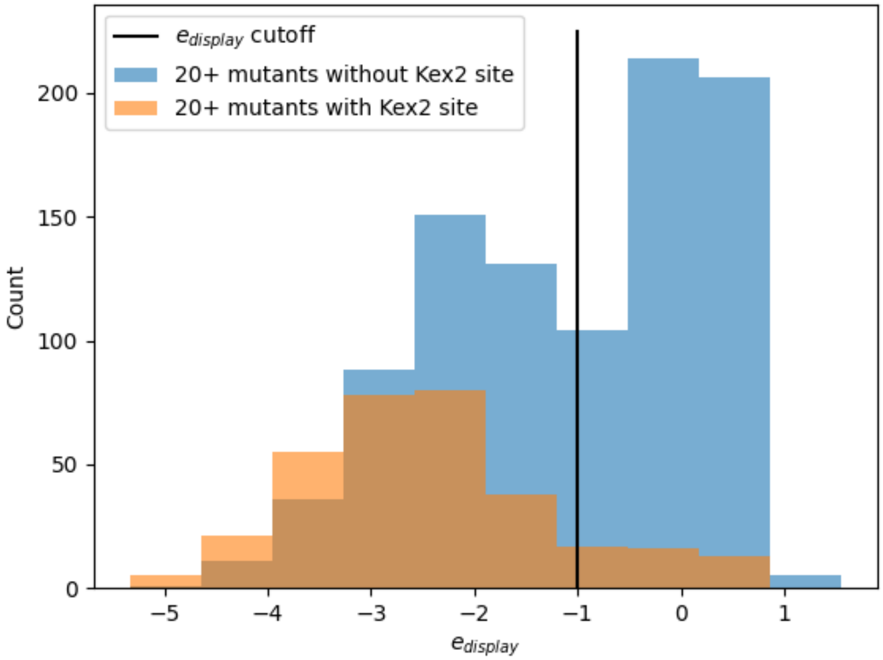


**Figure S7. 20+ mutants with Kex2 sites display less than 20+ mutants without Kex2 sites.** We examine $e_{disp}$ scores as they relate to presence or absence of Kex2 sites for 20+ mutants since designs with many mutations can be prone to unfolding. Kex2 can cleave unfolded proteins in the yeast endoplasmic reticulum if they have a Kex2 cleavage site. We classify sequences as having a Kex2 site if they contain the consensus sequences KR and RR[3]. Variants without Kex2 sites may still display or not display depending on other factors.

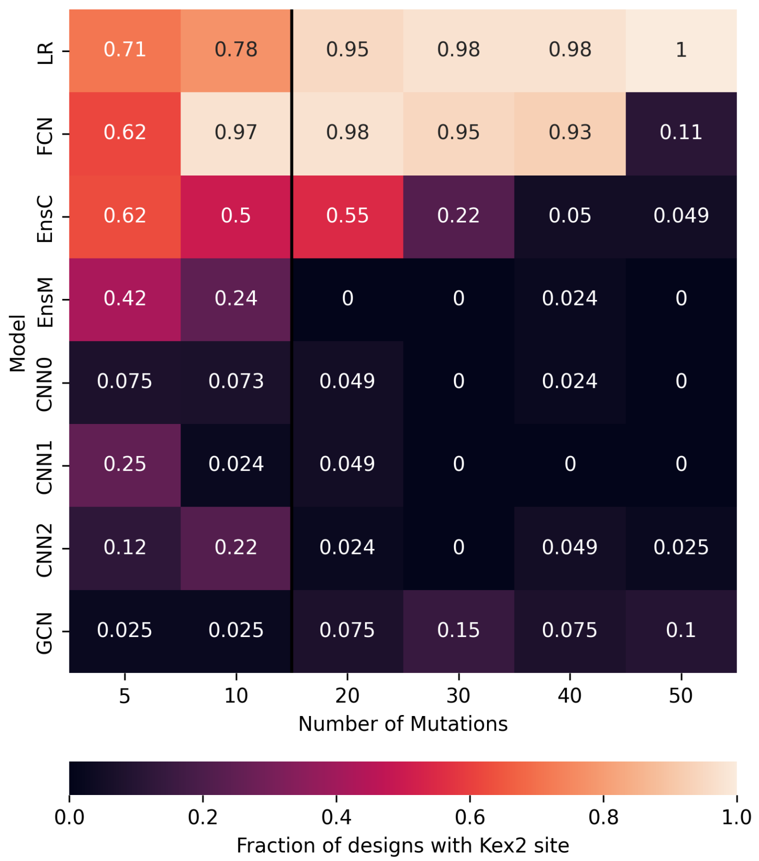


**Figure S8. LR and FCN produce higher proportions of designs with Kex2 sites.** We broadly determine possible Kex2 sites with the consensus sequences KR and RR and find the proportion of designs from each category with these consensus sequences. The black line separates the designs that are more likely to fold/not fold, based on the number of mutations, since folding hinders Kex2 cleavage even if a Kex2 cleavage site is present.

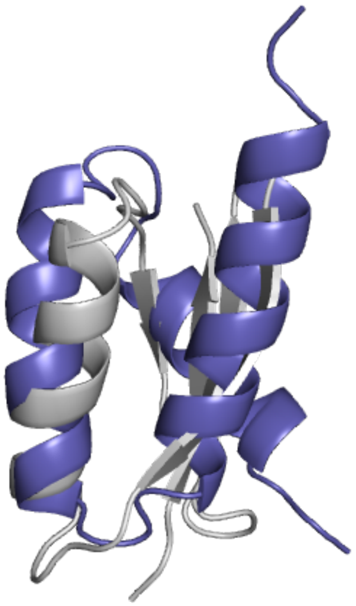

**Figure S9. GCN-40 is predicted to fold into a new topology distinct from GB1**. GCN-40, shown in purple, displays 2.3-fold higher than wildtype GB1, shown in grey, and is predicted by Alphafold to fold into a triple helix.

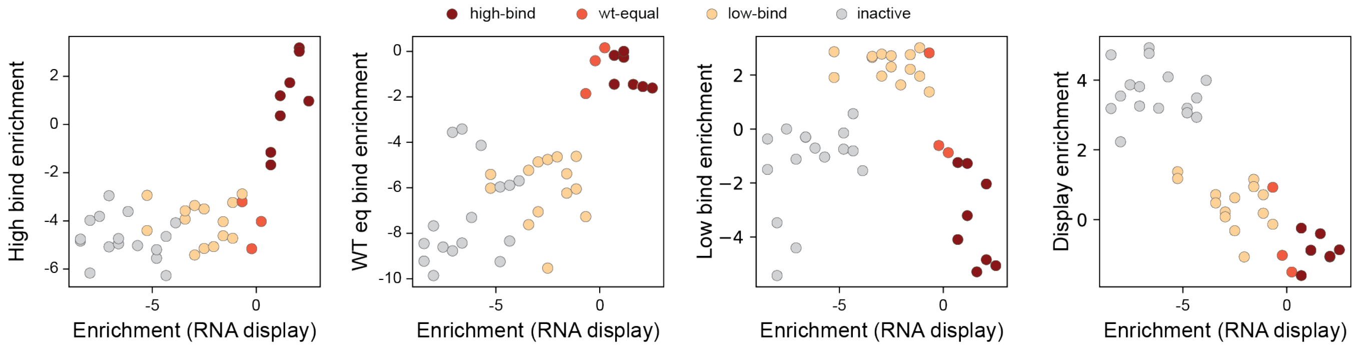


**Figure S10. Categorical binding characterization correlation with binding enrichment from RNA display.** To confirm that designs are placed in the correct category, we examine the enrichments of our 25 calibration sequences for each sorted population and for Olson et al.'s RNA display binding assay. We categorize each design as high-bind, wt-equal, low-bind, or inactive if the design has high enrichment in one of the populations, beyond our manually set threshold.
